# Avs2 drives non-canonical tetrameric assembly of trypsin-like domain for anti-phage defense

**DOI:** 10.64898/2026.08.25.746987

**Authors:** Lijie Guo, Pingping Huang, Wenhui Liu, Jingxian Liu, Dongyang Xu, Shengbin Yu, Zongqiang Wang, Lei Zhang, Zhaoxing Li, Xu Cao, Qian Yang, Mengjun Cheng, Nannan Wu, Meiling Lu, Lian-Wen Qi, Yibei Xiao, Meirong Chen

## Abstract

Bacteria have evolved diverse anti-phage defense systems, with the antiviral STAND family (Avs) representing one of the most diverse and widespread, encompassing at least 90 distinct families, yet only Avs3, Avs4, Avs5 and Avs7 have been well characterized. Here, we elucidated the molecular mechanism of Avs2-trypsin-MBL system, where phage terminase recognition by Avs2 triggers coupled activation of the protease and nuclease activities of trypsin-MBL. Cryo-EM structure of Avs2-trypsin-terminase and biochemical analysis reveal that the binding of terminase ATPase domain triggers the assembly of Avs2 into tetramer, with two unique ATP molecules bridging ATPase active-site recognition by TPR domain. This tetramerization drives the fused trypsin into an active C4-symmetric assembly, an architecture distinct from the conventional non-defense trypsin. Our study unravels the activation mechanism of Avs2-trypsin-MBL system, expanding our understanding on commonality and diversity of widespread Avs-mediated anti-phage immunity, alongside the structural and functional adaption of trypsin.

## Introduction

Owing to the continuous threat of phage infection, bacteria have evolved a diverse array of anti-phage defense systems, which function at various stages of phage life cycles to prevent phage propagation^[1–5]^. Through combination of a sensor and an effector domain, most defense systems efficiently sense phage invasion signals and rapidly activate protective responses to counter phage infection^[6–10]^.

Antiviral STAND superfamily (Avs), a prokaryotic counterpart of eukaryotic NOD-like receptors (NLRs) for pathogen recognition^[11–13]^, is present in approximately 4-5% of sequenced prokaryotic genomes, broadly distributed across bacterial and archaeal lineages^[14]^, suggesting that Avs-mediated immunity is evolutionarily widespread and efficient. Avs proteins typically comprise an N-terminal lethal effector domain, a central STAND ATPase domain, and a C-terminal sensory domain with tetratricopeptide repeats ^[11,14–17]^. Recognition of phage proteins triggers oligomerization of the STAND ATPase domain and activation of the effector domain, resulting in phage restriction. The Avs families exhibit extensive diversity, arising from the evolutionary relationships and divergent sensory architectures^[14,18]^. Over 90 distinct Avs families have been identified, constituting one of the most diversified family of immune systems. To date, Avs3^[14]^, Avs4^[14]^, Avs5^[15,16]^ and Avs7^[17]^ represent the best-characterized members of the Avs families, activated by the phage terminase, portal protein, JADA protein, and major capsid protein, respectively. Structural analysis of Avs families above has revealed how phage protein recognition by sensory domains is coupled to ligand-induced conformational changes and oligomerization of STAND ATPase domains, thereby activating downstream effector domains^[14,16,17]^. Compared with the above well-demonstrated Avs types, we have little knowledge on the Avs2 system, except that preliminary screening suggests that phage terminase could activate Avs2-mediated immunity. On the other hand, combination with a broad repertoire of effectors, such as nuclease, SIR2, and protease, further enriches the Avs families^[14–17]^. However, so far, most well-demonstrated Avs proteins are limited to those containing nuclease effector domains, either Mrr or Cap4 nuclease, which confer anti-phage immunity through nucleic acid degradation^[14,17]^. Since the structural and functional diversity of Avs families, it would be significant to uncover the novel effect mediated by Avs, such as protease or other enzymatic activities, and Avs proteins associated with non-nuclease effectors would fill the gap and broad our mechanistic understanding on these efficient and divergent families.

Our previous study has identified a widespread Avs2 group coupled with trypsin-MBL effector module, which confers bacteria strong anti-phage immunity^[19]^. In this study, we focused on the working mechanism of Avs2-trypsin-MBL. Biochemical and cell-based assays revealed that phage terminase protein triggers the coupled activation of the protease and nuclease activities of trypsin-MBL, which act in tandem to restrict phage infection. Using cryogenic electronic microscopy (cryo-EM), we obtained the structure of tetrameric Avs2-trypsin-terminase complex, presenting the active state of Avs2-trypsin. Combined with biochemical evidences, our results revealed that the ATPase domain of terminase alone is enough to activate Avs2-trypsin, wherein two ATP molecules synergistically contribute to terminase recognition by tetratricopeptide repeats enriched domain and C-terminal domain of Avs2-trypsin. Subsequently, four central STAND ATPase domains tetramerize, bringing trypsin domains close together with C4 symmetry. Importantly, the tetramerization of trypsin domains is prerequisite for their activation, which differs from most conventional trypsin proteins functioning in monomeric state^[20]^. Taken together, our study provides comprehensive mechanistic insights into an Avs2 system associated with a widespread trypsin-MBL module, which expands our current understanding of bacterial strategies for sensing and combating phage infection, as well as the functional and structural peculiarity of defense related trypsin proteins.

## Results

### Terminase-dependent activation of Avs2-trypsin-MBL confers anti-phage immunity

In our previous study, we have reported trypsin-MBL module associated with multiple defense systems, among which the Avs family was the most abundant. Furthermore, Avs2-trypsin-MBL system from *Vibrio cholerae* was validated conferred bacterial strong resistance to phages^[19]^. Avs2-trypsin-MBL system consists of two components: an Avs2 protein fused with an N-terminal effector that adopts a fold reminiscent of trypsin domain, and a metallo-β-lactamase (MBL) superfamily protein **(Fig.1A)**. The catalytic mutant of trypsin domain and MBL, trypsin^H32A^and MBL^D65A^, failed to defend against phage ΦCP-EC-23022 (a T4-like phage) **(Fig.1A)**, indicative of emzymatic activity of trypsin and MBL are essential for anti-phage function of Avs2-trypsin-MBL system.

**Fig. 1.**
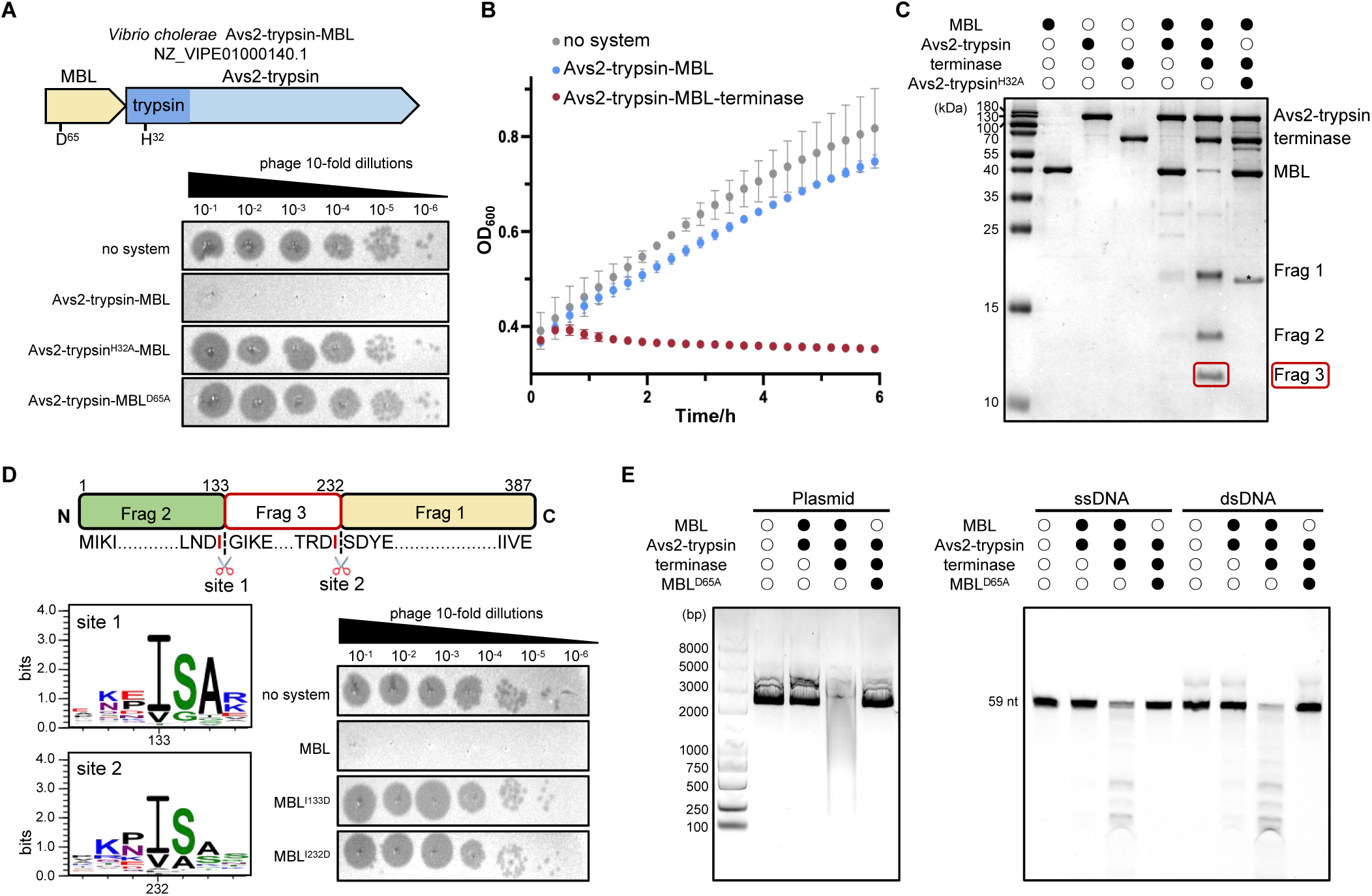
phage terminase activates Avs2-trypsin-MBL to function as an anti-phage defense system. (A) Phage plaque assays of phage ΦCP-EC-23022 on *E. coli* lawns expressing wild-type Avs2-trypsin-MBL or the catalytic site mutants. The schematic representation of *Vibrio cholerae* Avs2-trypsin-MBL operon annotated with predicted catalytic residues is shown on top. (B) Growth curves of *E. coli* expressing the Avs2-trypsin-MBL system with or without phage terminase, with a no-system strain as control. Curves are shown as the means ± SD of three independent biological replicates. (C) SDS-PAGE analysis of the MBL cleavage by Avs2-trypsin with or without terminase. The MBL fragments are labeled as Frag 1-3. A band of impurity is indicated with an asterisk. (D) Mapping of MBL cleavage sites by mass spectrometry. The Frag 3 from (C, lane 7) was sequenced. Cleavage sites are mapped onto the primary structure of MBL (top). The N- and C-terminal sequences of the Frag 3 are aligned below, with cleavage site isoleucine highlighted in red. The conservation analysis of the cleavage site among MBL from Avs2-trypsin-MBL systems is shown at the bottom. The importance of cleavage site was validated by Phage plaque assays (bottom right). (E) Agarose gel and urea-PAGE analysis of DNA cleavage by activated MBL.

It has been reported that that large terminase subunit, a key component of the phage DNA-packaging machinery, is recognized by Avs2 with various of N-terminal lethal effector domains^[14,21]^. To examine whether Avs2-trypsin-MBL can be activated by terminase, we performed cell toxicity assays where Avs2-trypsin-MBL and terminase from phage ΦCP-EC-23022 were co-expressed in *E. coli*. While *E. coli* expressing Avs2-trypsin-MBL alone grew normally as *E. coli* with no system, *E. coli* co*-*expressing Avs2-trypsin-MBL and terminase exhibit growth arrest **(Fig.1B)**, which indicates that terminase activates Avs2-trypsin-MBL to confer defense function.

Next, we purified Avs2-trypsin, MBL and terminase proteins to investigate the terminase-triggered activation of Avs2-trypsin-MBL in vitro. The result showed that while MBL incubated with Avs2-trypsin alone yielded little degradation, the addition of terminase led to significant cleavage of MBL into three specific fragments by Avs2-trypsin **(Fig.1C)**. The cleavage of MBL was abolished when the active site of trypsin domain was mutated **(Fig.1C)**. Mass spectrometry mapped the cleavage sites on MBL precisely after Ile133 and Ile232 **(Fig.1D and Supplementary Data)**. The residues isoleucine, followed by a small residue S/G/A, exhibited a high degree of sequence conservation in Avs2-trypsin-MBL systems across different strains **(Fig.1D)**. Mutation at either Ile133 or Ile232 resulted in a loss of antiviral activity of the Avs2-trypsin-MBL system **(Fig.1D)**. Consistently, this isoleucine-specific recognition was also observed in a subgroup of Hachiman-trypsin-MBL^[22]^, further suggesting a unique cleavage preference by trypsin domain in this trypsin-MBL module. Next, the activation of MBL by terminase-induced trypsin protelysis was further examined. Our previous study indicated that the Hachiman associated MBL protein showed robust DNase activity^[19]^. Thus, the Avs2-trypsin-MBL system incubated with or without terminase was subjected to DNase assay with various DNA substrates, including plasmids, ssDNA or dsDNA. The results showed that MBL protein or Avs2-trypsin-MBL alone was incapable of DNA cleavage, whereas MBL protein, but not the catalytic mutant, degraded all above DNA substrates only when the terminase protein was added to Avs2-trypsin-MBL **(Fig.1E)**.

We have reported that a unique loop insertion flanked by two proteolytic cleavage sites is responsible for maintaining MBL protein in an autoinhibited state and that proteolytic cleavage of this regulatory loop by trypsin domain releases the DNase activity of MBL^[19]^ **(Fig.S1A)**. To further visualize how this loop insertion affects MBL function, we tried crystallization of cleaved VcMBL^D65A^ from Avs2 system and KpMBL^D73A^ from Hachiman system, respectively. We obtained high-quality crystal of cleaved KpMBL^D73A^ and determined the structure at a resolution of 2.7 Å, which reveals an overall typical MBL fold **(Fig.S1B and Supplementary Table 1)**. After cleavage, the regions (108–128 aa and 189–196 aa) near the two cleavage sites on the loop insertion was not resolved in the structure **(Fig.S1B)**, indicating its substantial conformational flexibility after cleavage. With the two regions released by two-site cleavage, a basic channel at MBL surface with the active site cavity binding Zn^2+^ is exposed **(Fig.S1B)**, shaped to make sufficient room for accommodating substrate DNA.

Taken together, these findings suggest that upon phage infection, the phage terminase activates Avs2-trypsin to release its protease activity, which cleaves the inhibitory loop insertion of MBL and converts it into active form for DNA degradation, resulting in anti-phage immunity through abortive infection.

### Phage terminase induces Avs2-trypsin forming unique tetramer

To elucidate how Avs2-trypsin is activated by phage terminase, we firstly investigated their interaction in vitro by gel filtration. While Avs2 alone exists as monomer, the incubation of Avs2-trypsin with terminase leads to significant shift of the peak **(Fig. 2A)**, indicative of a higher-order oligomerization complex formation of Avs2-trypsin with terminase. Then, we applied this Avs2-trypsin-terminase complex in the presence of ATP and Mg^2+^ onto cryo-EM to determine the active state of Avs2-trypsin. 2D classification yields well-defined class averages displaying tetrameric state of the complex **(Fig. 2B)**. Further 3D construction and refinement result in a map of Avs2-trypsin-terminase at a resolution of 2.5 Å **(Fig. S2, S3 and Supplementary Table 2)**. Avs2-trypsin-terminase exists as a C4-symmetric tetramer, with each Avs2-trypsin protomer binding one terminase **(Fig. 2C)**.

**Fig. 2.**
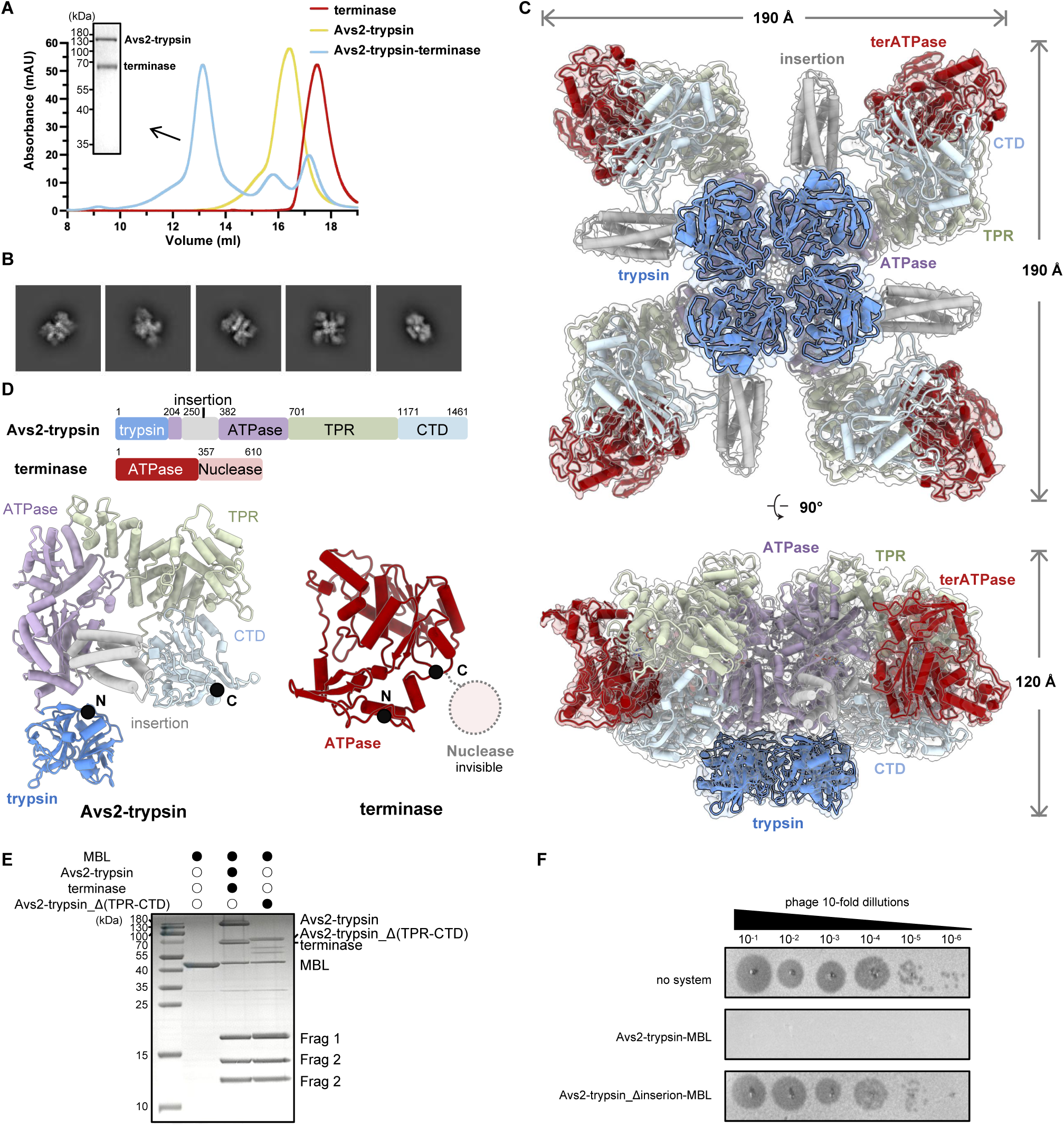
Phage terminase induces tetramerization of Avs2-trypsin. (A) Comparative size-exclusion chromatography profiles of Avs2-trypsin, phage terminase and reconstituted Avs2-trypsin-terminase complex. The peak of the complex was analyzed by SDS-PAGE. (B) Representative 2D class averages of Avs2-trypsin-terminase complex. (C) Cryo-EM structure of the Avs2-trypsin-terminase complex shown in two orthogonal views. The structure model is shown in cartoon with cryo-EM map. The trypsin domain, insertion domain, STAND ATPase domain, tetratricopeptide repeats enriched domain (TPR) and C-terminal domain (CTD) of Avs2-trypsin protomer are colored in blue, gray, purple, palegreen and palecyan, respectively, with the ATPase domain of terminase (terATPase) colored in red. Each Avs2-trypsin protomer binds one terminase subunit. (D) The cartoon representation of Avs2-trypsin protomer and terATPase. The unresolved nuclease domain of terminase (terNuclease) is indicated with a pink dashed-line circle. The N- and C-terminus are indicated with black spheres. The primary structures of Avs2-trypsin and phage terminase are shown on top. (E) SDS-PAGE analysis of MBL cleavage by Avs2-trypsin_Δ(TPR-CTD). (F) Phage plaque assays of phage ΦCP-EC-23022 on *E. coli* lawns expressing wild-type Avs2-trypsin-MBL or Avs2-trypsin_Δinsertion-MBL.

The Avs2-trypsin protomer consists of a N-terminal trypsin domian (1–203 aa), a central STAND ATPase domain (204–700 aa) with a unique insertion (250–381 aa), a tetratricopeptide repeats enriched domain (TPR, 701–1170 aa) and a C-terminal domain (CTD, 1171–1461 aa). For the structure of the terminase, while the C-terminal nuclease domain (terNuclease, 357–610 aa) is mostly invisible, the N-terminal ATPase domain (terATPase, 1–356 aa) adopts a typical ASCE (additional strand, conserved E) ATPase fold with a Rossmann-like α/β architecture, comprising a central six-stranded parallel β-sheet surrounded by α-helices^[23–25]^ **(Fig. 2D)**. In the Avs2-trypsin-terminase tetramer, the terATPase binds each Avs2-trypsin protomer in between TPR and CTD. Four Avs2-trypsin-terminase oligomerize at central STAND ATPase domains into symmetric tetramer. The tetramerization of STAND ATPase domain drives the fused N-terminal trypsin domais into close proximity with C4 symmetry **(Fig. 2C)**.

The TPR and CTD in Avs families are believed to function as sensory architectures to recognize the pattern of invasive signals. However, whether it’s indispensable for the activity of Avs families has not been explored. In this study, we surprisingly found that deletion of TPR and CTD results in automatically activation of Avs2-trypsin even in the absence of terminase **(Fig. 2E)**, suggesting its regulatory role in not only activation but also the auto-inhibition of Avs systems to precisely control anti-phage immunity.

The STAND ATPase domain scaffolds tetrameric assembly of Avs2-trypsin. Besides the typical the nucleotide-binding (NBD), helical domain 1 (HD1) and winged-helix (WHD) subdomains, the STAND ATPase domain of Avs2-trypsin harbors a unique α-helix-rich insertion, which is absent in all other reported Avs types **(Fig. S4, left)**. Deletion of this insertion region leads to completely loss of anti-phage activity of Avs2-trypsin-MBL system **(Fig. 2F)**, suggesting its important role in Avs2 function and distinguishing Avs2 from other Avs types with canonical STAND ATPase domains. Akin to other types of Avs proteins, an ATP molecule coordinated with Mg^2+^ is embedded in the NBD subdomain, with the phosphates and ribose recognized by walker A involving T392, G393, K394, S395, H396 and R499 **(Fig. S4, right)**, mediating the subsequent activation of downstream trypsin effector^[14]^.

Unlike previously reported Avs proteins that mostly adopt a tetrameric but stretched conformation, Avs2-trypsin-terminase complex exhibits an overall flat architecture **(Fig. S5A)**. Specially, compared to Avs3^[14]^, TPR in Avs2 possesses less α-helix repeats and leaves larger space for terNuclease **(Fig. S5B)**, which may account for the flexibility and invisibility of terNuclease in the structure of Avs2-trypsin-terminase.

### Two ATP molecules mediate terminase recognition by Avs2-trypsin

The recognition of terminase by Avs2-trypsin is distinct with Avs3^[14]^. In the Avs2-trypsin-terminase structure, the terATPase is shape-complementarily embedded in the cavity formed by TPR and CTD of Avs2-trypsin. By contrast, only weak and fragmented density of terNuclease can be observed in the cavity, preventing the reliable building of an atomic model **(Fig. 3A and S6)**. Since the terNuclease is not rigidly fixed and no obvious contact with Avs2-trypsin could not be observed, we guessed that the terNuclease is not required and terATPase alone is sufficient to activate the Avs2-trypsin. To test the hypothesis, we purified isolated terATPase and terNuclease individually and assessed their ability to activate Avs2-trypsin. The results showed that terATPase alone, but not terNuclease, was able to induce the oligomerization of Avs2-trypsin and exhibited comparable abilities to full-length terminase in activating Avs2-trypsin to cleave MBL **(Fig. 3B, 3C)**. These results indicate that the terATPase is sufficient to activate Avs2-trypsin, suggesting that Avs2-trypsin has a relaxed requirement for terminase recognition.

**Fig. 3.**
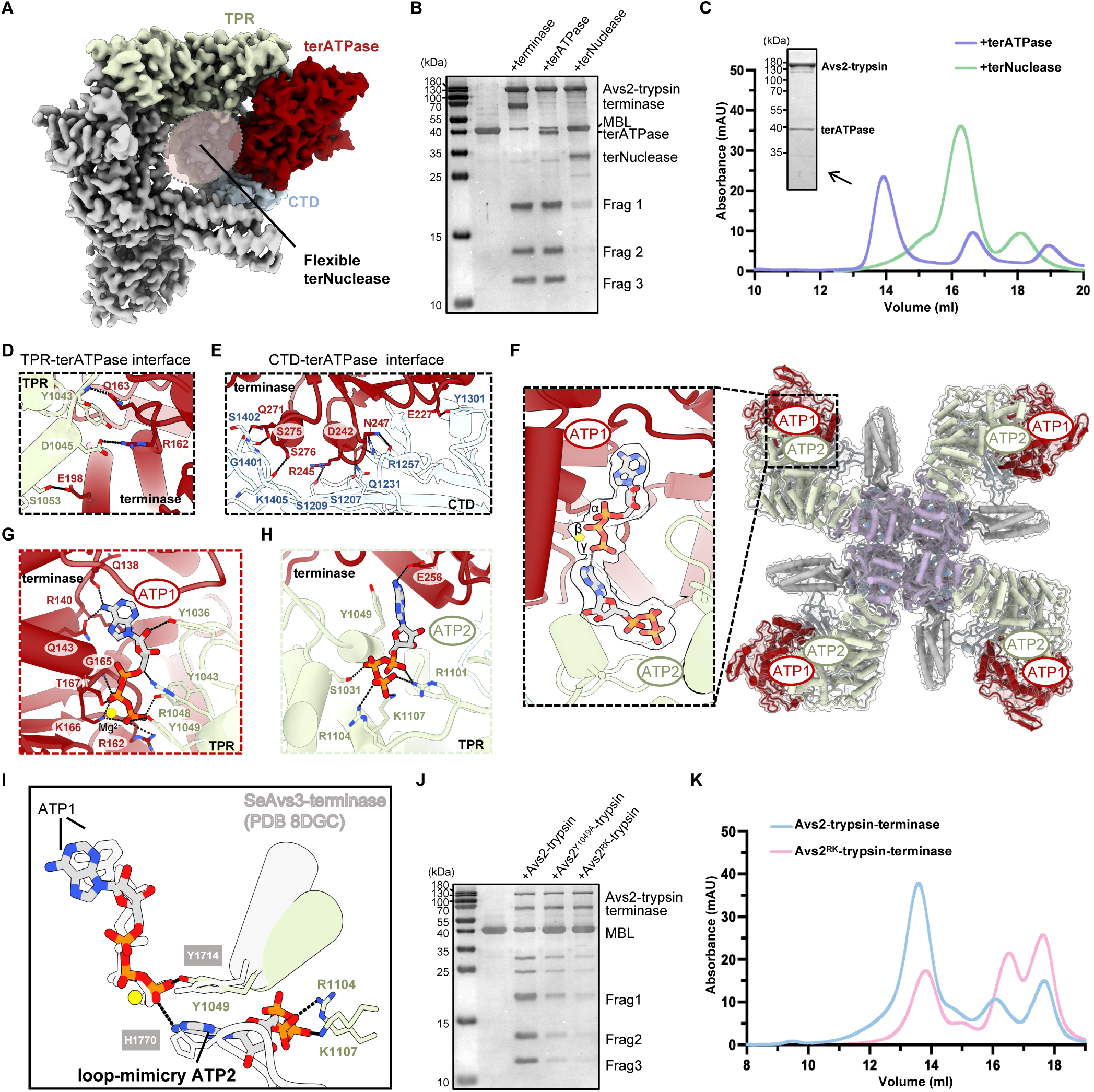
Structural basis of terminase recognition by Avs2-trypsin. (A) The close-up view of terATPase embedded in cleft formed by TRP (palegreen) and CTD (palecyan). The position of the unresolved terNuclease is indicated with a pink dashed circle. The Avs2-trypsin-terminase protomer is shown as density map. (B) SDS-PAGE analysis of MBL cleavage by Avs2-trypsin in the presence of terATPase or terNuclease. (C) Size-exclusion chromatography analysis of the complex formation of Avs2-trypsin with terATPase or terNuclease. The peak of complex was examined by SDS-PAGE. (D, E) Close-up views of the interface TPR-terATPase (D) and CTD-terATPase (E). The residues involved in the interactions are labeled. (F) Two unique ATP molecules (ATP1 and ATP2) bound in each Avs2-trypsin protomer. The inset shows the coordination of ATP1 and ATP2. (G, H) Close-up views of ATP1 (G) and ATP2 (H) recognition by terATPase and TPR. (I) Structure superimposition of Avs2-trypsin-terminase onto *Se*Avs3-terminase shows the overlapping of ATP2 and residue H1770 of *Se*Avs3. Avs2-trypsin is in color and *Se*Avs3 in light grey. (J) SDS-PAGE analysis of MBL cleavage by the mutants Avs2-trypsin^Y1049A^ and Avs2^RK^-trypsin. RK: mutant with R1104A/K1107A. (K) Size-exclusion chromatography analysis of complex formation of Avs2^RK^-trypsin with terminase.

terATPase is extensively recognized by TPR and CTD of Avs2-trypsin via a network of polar and electrostatic interactions. Specifically, at the interface between Avs2-trypsin TPR and terATPase, Y1043, D1045, and S1053 from Avs2-trypsin TPR establish hydrogen bonds with Q163, R162, and E198 from terATPase, respectively **(Fig. 3D)**. The CTD and terATPase form a broad and complementary interface, which is maintained by residues S1207, S1209, R1257, Q1231, Y1301, G1401, S1402 and K1405 from CTD together with E227, D242, R245, N247, Q271, S275 and S276 from terATPase **(Fig. 3E)**.

An ATP molecule (ATP1) located in the catalytic pocket of terATPase further reinforces this recognition through direct interactions with both terATPase and TPR **(Fig. 3F and 3G)**, which is similar to ATP1 in *Se*Avs3^[14]^. Within terATPase, G165/K166/T167 (walker A motif) and R162 form hydrogen bonds with the β-and γ-phosphate of ATP1, whereas Q138, R140, and Q143 stabilize the ribose moiety **(Fig. 3G).** In TPR, three phosphates groups of ATP1 contact with Y1036, Y1043, R1048, and Y1049 **(Fig. 3G)**. Through these interactions, ATP1 bridges the recognition between terminase and TPR in Avs2-trypsin-terminase complex.

Strikingly, near ATP1, an additional ATP molecule (ATP2) was observed, with its ribose moiety forming hydrogen bonds with the γ-phosphate of ATP1 **(Fig. 3F)**. Sandwiched by TPR and terATPase, ATP2 extensively interacts with TPR mediated by S1031, R1101, R1104, K1107 and Y1049, and establishes hydrogen bond with E256, the conserved catalytic residue of terATPase **(Fig. 3H)**.

Given that ATP2 specifically contacts ATP1, Avs2-trypsin, and terminase, it is likely functionally relevant rather than incidental. To test this hypothesis, we superimposed the Avs2-trypsin-terminase structure onto that of SeAvs3-terminase, which contains ATP1 but lacks ATP2. In this alignment, ATP1 and its recognition residue (Y1049 in Avs2-trypsin versus Y1714 in SeAvs3) are well matched in the two structures. However, close inspection reveals that the H1770-containing loop, which is essential for ATP1 binding and thus indispensable for Avs3 immunity^[14]^, is entirely absent in Avs2-trypsin. Instead, ATP2 occupies the corresponding position, with its purine moiety aligning with H1770 and forming a π–π stacking interaction with Y1049, while the phosphate groups are stabilized by R1104 and K1107 **(Fig. 3I)**. These observations suggest that ATP2 in Avs2-terminase probably mimics H1770 in SeAvs3-terminase, to strengthen recognition between Avs2 and terATPase via ATP1 binding. Supporting our idea, all the mutations of the involving residues, Y1049A, R1104A and K1107A, would weaken the activation of Avs2-trypsin by terminase for MBL cleavage **(Fig. 3J)**. Gel filtration analysis further confirmed that this impaired function results from the declined tetramerization tend of Avs2-trypsin-terminase **(Fig. 3K)**, owing to the diminished interactions between ATP1/2 and the mutant. These results together suggest that both ATP1 and the loop-mimicry ATP2 contribute to the activation of Avs2 via mediating recognition of terminase.

Collectively, the above data reveals the infection recognition mode of Avs2-trypsin and the unique structure elements contributing to phage terminase recognition, where two essential ATP molecules synergistically mediate the recognition and activation of Avs2-trypsin.

### Trypsin domains adopt non-canonical tetrameric assembly for defense function

We previously reported the structure of active HamAB-trypsin bound with the trigger DNA, revealing that the oligomerization of sensory protein HamAB is essential for activation of trypsin domain^[19]^. However, the trypsin domain is unresolved in that structure, leaving a question that in trypsin-MBL associated defense how the sensory protein activates trypsin domain. In this study, the structure of Avs2-trypsin-terminase unveils that driven by tetramerization of central STAND ATPase domains, four well defined N-terminal trypsin domains form a tetramer with an identical C4 symmetry as the fused STAND ATPase domains, distinct from the C2-symmetric arrangement adopted by the effector domains of other Avs families^[14,17]^ **(Fig. 4A and S5A)**. The structure of Avs2-trypsin-terminase, for the first time, reveals the conformation of active trypsin domain in trypsin-MBL associated defense system. Since most known conventional trypsin proteins function as monomer, the working mechanism of this non-canonical assembly of trypsin domains raises our interests.

**Fig. 4.**
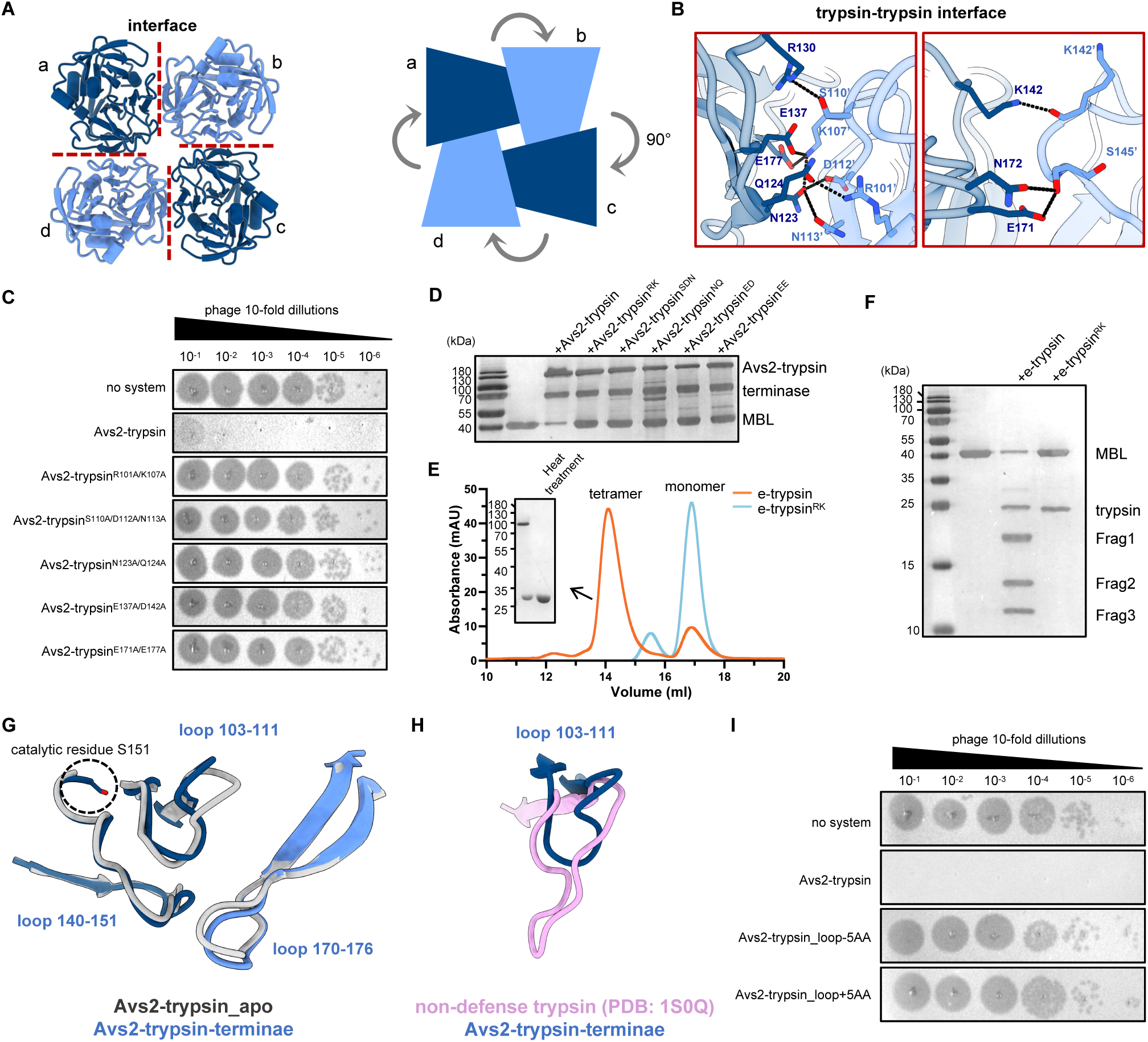
Tetramerization of trypsin is required for defense function. (A) The C4-symmetric assembly of trypsin domains. The four subunits are assigned as a-d. Two interaction regions at trypsin–trypsin interface are indicated with the black and red dashed lines. (B) Detailed views of trypsin-trypsin interface. The residues involved in the interaction are labeled. (C) Phage plaque assays of phage ΦCP-EC-23022 on *E. coli* lawns expressing trypsin interface mutants. (D) SDS-PAGE analysis of MBL cleavage by trypsin interface mutants in the presence of terminase. (E) Size exclusion chromatography analysis of tetramerization of e-trypsin domain and e-trypsin-based trypsin–trypsin interface mutant (e-trypsin^R101A/K107A^). e-trypsin refers to trypsin variant with tetramerization capacity enhanced, which was engineered by mutating the aspartate residue D108 that mediates electrostatic repulsion. (F) SDS-PAGE analysis of MBL cleavage by interface mutant e-trypsin^R101A/K107A^. (G) Structure superimposition of the Avs2-trypsin in apo form onto that in active state reveals the conformational changes of three loops at trypsin-trypsin interface. The apo form Avs2-trypsin is predicted by AlphaFold 3 with a pTM score of 0.8. (H) Structural alignment of the loop 103–111 in the trypsin domain of Avs2-trypsin-terminase with the corresponding region in non-defense monomeric trypsin protein (PDB: 1S0Q). (I) Phage plaque assays of phage ΦCP-EC-23022 on *E. coli* lawns expressing Avs2-trypsin mutant carrying shortened or extended loop 103–111. Avs2-trypsin_loop-5AA and Avs2-trypsin_loop+5AA refer to the mutants of loop 103– 111 with 5 animo acids truncation and extension, respectively.

To elucidate the relationship between tetramerization and activation of trypsin domain, we first conducted a comprehensive analysis of tetrameric interface of trypsin domains. In trypsin tetramer, burying a surface area of 2564 Å² in total, two adjacent two trypsin domains contact at same interface through extensive hydrogen bonds, involving residues N123, Q124, R130, E137, K142, E171, N172 and E177 from one trypsin domain and R101’, K107’, S110’, D112’, N113’, K142’ and S145’ from another **(Fig. 4A and 4B)**. The importance of these interactions on MBL cleavage as well as Avs2-trypsin-MBL defense function were examined by mutagenesis assay. As a result, all the trypsin mutants failed to cleave MBL protein, and completely abolished anti-phage immunity **(Fig. 4C, 4D)**. Due to strong tetramerization ability of STAND ATPase domain in Avs2-trypsin, it cannot be directly concluded that trypsin domain with above interface mutations dissociates in full-length Avs2-trypsin and the loss of function of trypsin domain is resulted from the disruption of trypsin tetramer. As trypsin domain alone hardly forms tetramer under gel filtration condition, to confirm the trypsin tetramer mediated activation, we screened and obtained an enhanced mutant of trypsin domain, trypsin^D118S/N119S^ (e-trypsin) **(Fig. S7A)**. The e-trypsin displays strong tetramerization ability in gel filtration and retains robust protease activity **(Fig. S7B and S7C)**, which can be used to characterize the effects of interface residues on trypsin tetramerization formation. As a result, the e-trypsin with mutation R101A and K107A cannot form tetramer in gel filtration and failed to cleave MBL **(Fig. 4E and 4F)**, suggesting that the disruption of trypsin-trypsin interface leads to its disassociation, thus impairing protease activity. Based on the results above, we concluded that tetramerization of trypsin domains is indispensable for its protease activity and defense function.

We next investigated how the tetramerization activates trypsin domain. Structural comparison of the monomeric and the active tetrameric Avs2-trypsin revealed conformational shifts of three closely associated loops at trypsin-trypsin interface, including loop 103–111, loop 140–151 and loop 170–176 **(Fig. 4G)**. Particularly, the loop 140–151 has close proximity to the catalytic site **(Fig. 4G)**. Importantly, the loop 103–111 is highly conserved in trypsin-MBL defense module **(Fig.S8)**, but distinct with that of non-defense trypsin proteins **(Fig.4G)**. Structural alignment of trypsin from Avs2 with other non-defense monomeric trypsin proteins reveal significant difference in this loop, as the latter possesses longer loop extended with about 5 amino acids **(Fig.4H)**. To validate the importance of this loop for trypsin activation in Avs2-trypsin, we assessed the anti-phage activity of Avs2-trypsin with the loop shortened or extended. The results showed that either truncation or extension of the loop with 5 amino acids would completely abolish the anti-phage function of Avs2-trypsin-MBL **(Fig. 4I)**, suggesting that this interfacial loop is precisely adapted to mediate the oligomerization-induced trypsin activation. The loop 103–111 further associates with the interfacial loop 140–151 and 170–176, potentially transferring the activation signal to the active site in proximity.

In summary, our results indicate that unlike canonical trypsin, defense related trypsin domain adopts a unique tetrameric architecture **(Fig. S9)**, where the interface loops mediate the activation of trypsin.

## Discussion

Combined with structural and biochemical results, we elucidated the molecular mechanism underlying Avs2-trypsin-MBL mediated anti-phage immunity: upon infection, Avs2-trypsin recognizes the ATPase domain of phage-encoded large terminase subunit, which, mediated by three ATP molecules, induces the tetramerization of Avs2-trypsin via assembly of STAND ATPase domains. Consequently, four trypsin domains are brought into close proximity to form non-canonical tetrameric architecture, which specifically cleaves and activates MBL to degrade DNA, thereby preventing phage propagation **(Fig. 5)**.

**Fig. 5.**
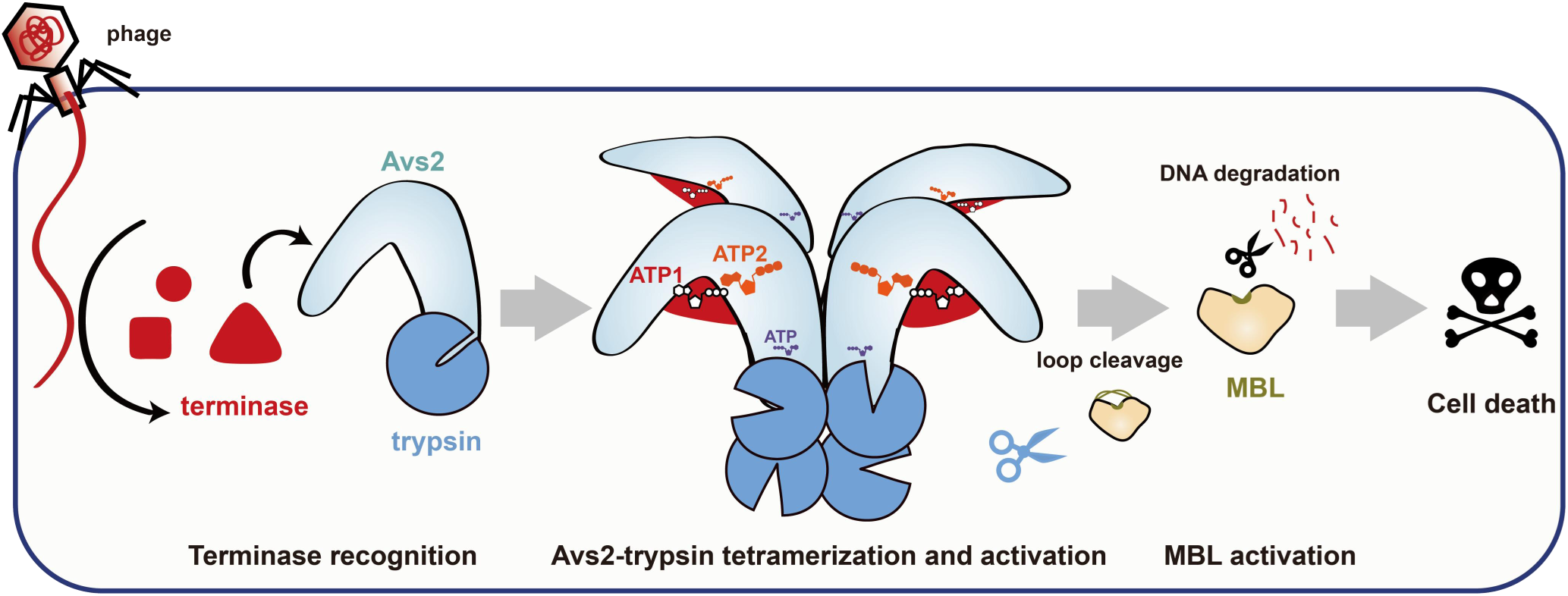
The proposed mechanism of Avs2-trypsin-MBL mediated anti-phage defense. Upon infection, Avs2-trypsin targets the ATPase domain of the phage-encoded large terminase subunit. This recognition event, facilitated by three ATP molecules, triggers the assembly of the STAND ATPase domain and drives Avs2 tetramerization. This structural rearrangement brings four trypsin domains into close proximity, forming a non-canonical tetrameric architecture that specifically cleaves and activates MBL. The activated MBL subsequently degrades DNA and thus lead to host death, effectively blocking phage propagation.

Avs families have evolved divergent molecular mechanisms for target recognition and immune activation. Avs1 and Avs3 recognize the phage large terminase subunit; Avs4 and Avs11 target phage portal protein; Avs7, Avs8 and Avs10 sense the major capsid protein; other Avs families further expand this recognition repertoire to include diverse phage proteins, such as tail tube, portal adaptor, and tail assembly chaperone^[14,17]^. Avs2-trypsin also recognizes the terminase, but employs a recognition mechanism distinct from that previously characterized for *Se*Avs3. While the activation of *Se*Avs3 requires recognition of both ATPase and nuclease domains of terminase^[14]^, terminase ATPase domain alone is sufficient to activate Avs2-trypsin. We proposed that the distinct recognition modes of Avs2-trypsin and *Se*Avs3 arose from differences in TPR length. The shorter TPR of Avs2-trypsin provides excessively large enclosure of the terminase, resulting in a more flexible nuclease domain configuration, whereas the longer TPR of *Se*Avs3 forms a dual-cavity architecture that engages both ATPase and nuclease domains. Notably, Avs2-trypsin employs a unique loop-mimicry ATP molecule to capture terminase, representing a novel target recognition pattern among the reported Avs families. Besides the pathogen recognition, the effector assembly further differentiates Avs2-trypsin from the well-studied nuclease-containing Avs systems in effector activation: while in Avs3, Avs4 and Avs7, the nuclease cap4 and Mrr mediate tetramerization by forming a C2-symmetric arrangement with the two dimers independently forming active sites, in Avs2-trypsin, trypsin domain adopts a C4 symmetry with tetramerization indispensable for activity. The structure of Avs2-trypsin could be well superimposed onto a recent reported Avs2 homolog. However, the effector domain of this Avs2 homolog remains functionally uncharacterized and structurally unresolved^[26]^**(Fig. S5A).** Collectively, target recognition and downstream effector activation can be modularly and independently diversified within the Avs families.

The structural role of the loop-mimicry ATP reminds us of Rad50-Mre11 complex, wherein ATP is sandwiched between two Rad50 head domains and directly contributes to the dimer formation^[27,28]^. Besides Rad50–Mre11, ABC transporter^[29]^ and SMC^[30]^ exhibit similar ATP-dependent dimerization mechanisms, highlighting ATP-mediated interface assembly. Beyond ATP, other nucleotides can also contribute to protein assembly by promoting or stabilizing intermolecular interactions, such as GTP in Tubulin^[31]^ or FtsZ^[32]^, c-di-GMP in PilZ^[33]^ or LapD^[34]^, and cGAMP in STING^[35]^, presenting an alternative strategy for regulating complex association. Additionally, the function of loop-mimicry ATP in Avs2-trypsin-terminase may enable precise recognition of phage infection and Avs2-trypsin activation under stringent control of cellular ATP level to avoid mis- or over-activation.

Our results showed that tetramerization of defense-related trypsin domain is required for releasing its protease activity. To our knowledge, most previously characterized trypsin-like serine proteases function in monomeric state after activation through proteolytic removal of the N-terminal peptide, which generates a new N terminus that stabilizes the active-site architecture through formation of the conserved Ile16-Asp194 salt bridge^[36–39]^. Interestingly, human β-tryptase with destabilized salt bridge represents a notable exception, in which its proteolysis activity depends on tetramer assembly and stabilization of the active site ^[40,41]^ **(Fig. S9)**. By contrast, although the defense-related trypsin domain also adopts tetramer architecture, with residues corresponding to Ile16 and Asp194 absent, it presents a distinct activation mode, where the tetramerization activates trypsin domains through conformational shifts of the interfacial active-site-proximal loops. Notably, the loop 103–111 of trypsin domain has its structural counterpart in human β-tryptase, the 147-loop, which is responsible for mechanical interlocking of tetramer for β-tryptase activation^[40,42]^. This structural similarity implies that the importance of the loop 103–111 for tetramerization and activation of trypsin domain. Interestingly, by means of AF3-based structure prediction, a preprint by Evans *et al*. proposed an alternative explanation for trypsin tetramer requirement, where tetramer assembly of trypsin may allow the simultaneous capture of two MBL cleavage sites for efficient activation^[43]^. However, this could not explain our observation that the disruption of tetramer completely abolished other than slowed down cleavage, supporting the indispensable role of trypsin tetramerization for active site shaping.

## Data availability

The atomic coordinates and structure factors for the crystal structure determined in this study have been deposited in the Protein Data Bank under accession code 44AF (Cleaved-loop MBL). Atomic coordinates, maps and structure factors of the reported cryo-EM structure have been deposited to the Protein Data Bank under accession number 27UW (Avs2-trypsin-terminase tetramer) and the EM Data Bank under accession codes EMD-81210, EMD-81216, EMD-81451, EMD-81453, EMD-81456 and EMD-81457.

## Acknowledgments

We would like to thank the Instrument Analysis Center (IAC) at Shanghai Jiao Tong University for Cryo-EM data collection. This work is supported by National Natural Science Foundation of China (32271330, 32471316, to M.C.; 82473977, 82321005, to Y.X.; 32470145, to N.W.; 82304614, to Z.L.; 32000441, to D.X.; 82373892, to X.C.), Basic Research Program of Jiangsu (BK20250200, to M.C.; SBK2024010634, to Y.X.), National Key Research and Development Program of China (2023YFC3402300, to M. L. and M.C.), STI2030-Major Projects (2021ZD0203400, to Y. X.), the Fundamental Research Funds for the Central Universities (2632025TD02, to M.C.; ZQN-924, to D.X.; 2632023GR17, to Z.L.).

## Author contributions

M.R.C., Y.B.X. and L.W.Q. supervised the project. L.J.G., P.P.H., W.H.L., J.X.L. and S.B.Y. performed cell-based assays, protein purification, and experiments. P.P.H., and M.R.C. determined the Cryo-EM structures. M.R.C., Y.B.X., L.W.Q., L.J.G., P.P.H., W.H.L., D.Y.X., J.X.L., S.B.Y., X.C., Z.Q.W., L.Z., Z.X.L., Q.Y., M.J.C., N.N.W. and M.L.L. analyzed data. L.J.G. and M.R.C. wrote the manuscript. All authors discussed the results and contributed to the final manuscript.

## Supplementary Figure legends

**Fig. S1.**
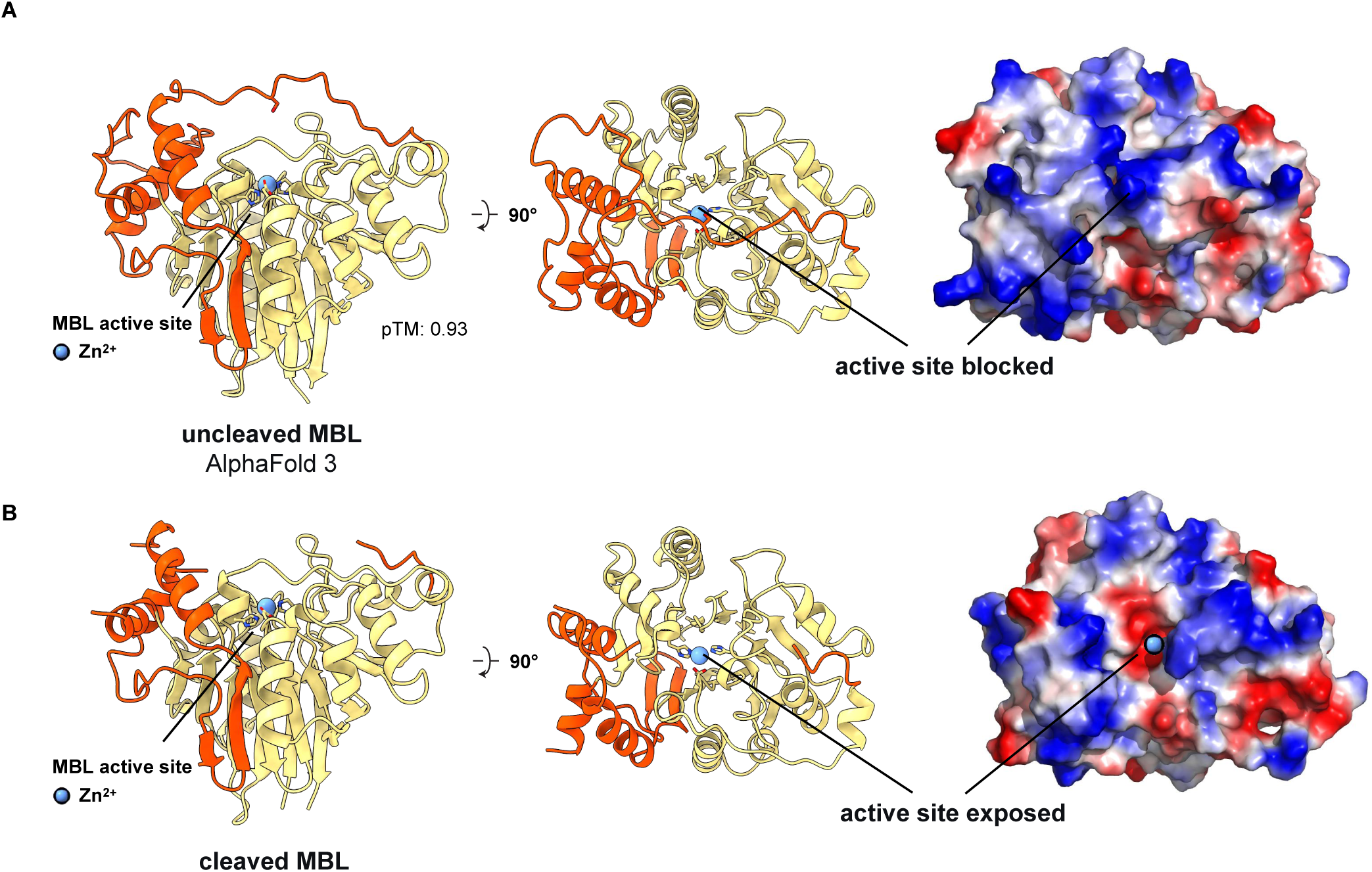
Structure comparison of MBL in uncleaved and cleaved states. (A) AF3-predicted structure model of uncleaved MBL presented in cartoon in front and top views, with a unique loop insertion colored in orange red covering the active site (left and middle). The electrostatic surface potential of the uncleaved MBL with the active site blocked (right). Zn^2+^ colored in blue is located in active site of MBL. (B) The crystal structure of the cleaved MBL presented in cartoon in front and top views, highlighting the exposed active site bound with Zn^2+^ (left and middle). The electrostatic surface potential of the cleaved MBL with the active site exposed (right).

**Fig. S2.**
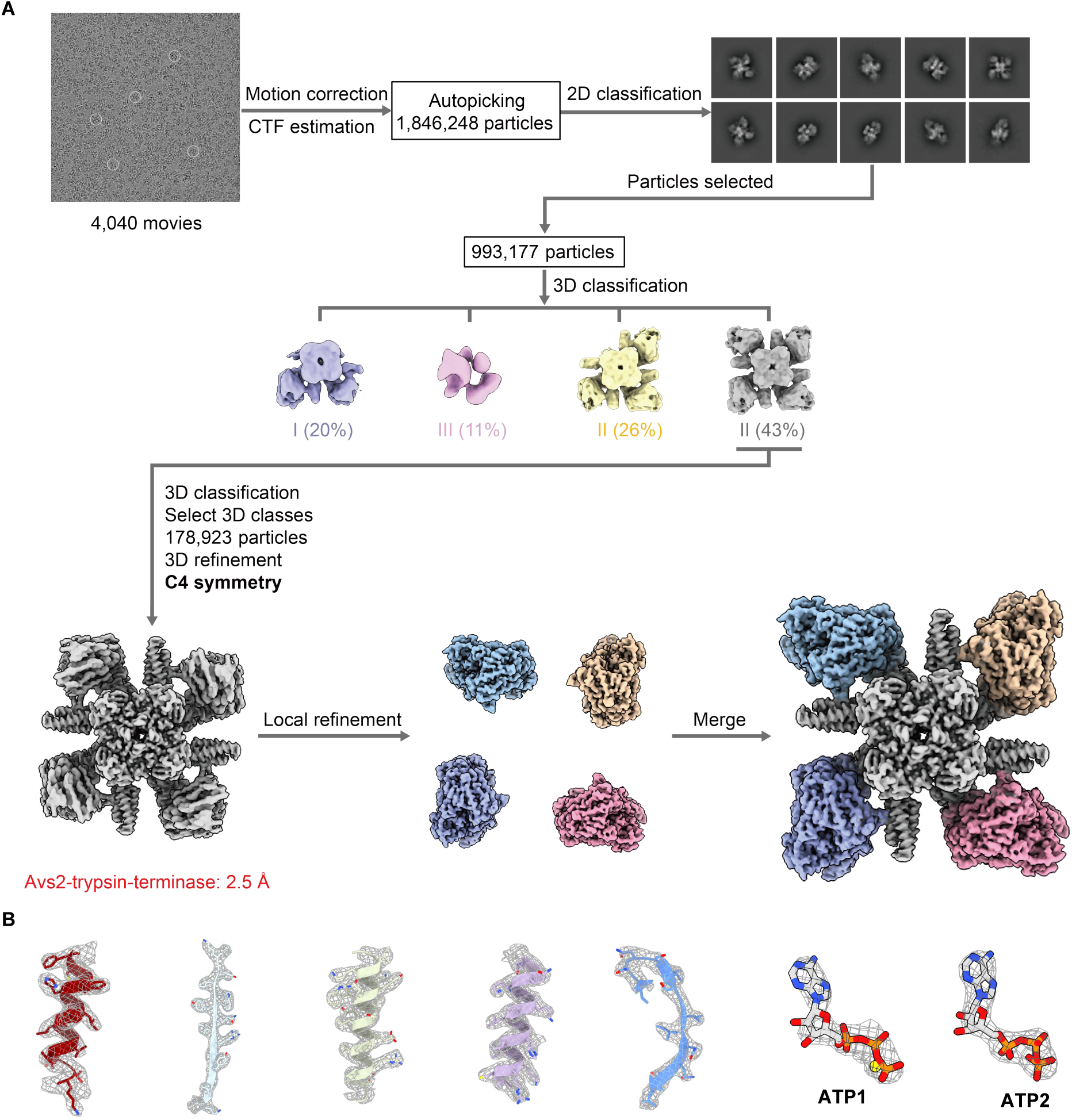
Single-particle cryo-EM analysis of Avs2-trypsin-terminase. (A) The flow-chat of single-particle cryo-EM analysis. (B) The representative density maps for selected regions from each domain and ATP1/2 molecules of Avs2-trypsin-terminase.

**Fig. S3.**
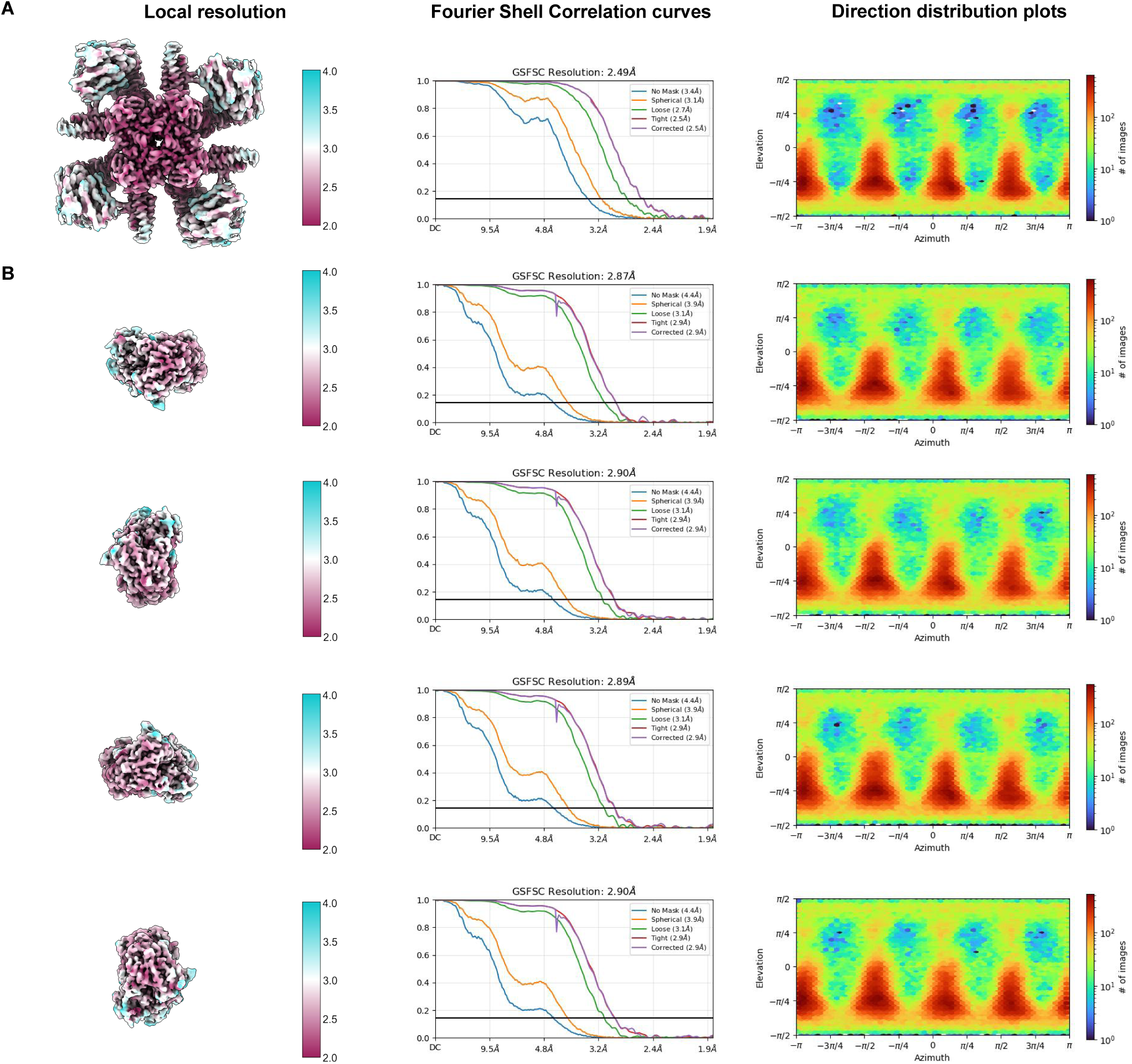
Cryo-EM density maps of the Avs2-trypsin-terminase complex. (A) Cryo-EM density map of the intact Avs2-trypsin-terminase colored by local resolution (left). The gold-standard FSC of 0.143 resolution graph and the viewing direction distribution plot are shown at middle and right. (B) The density map of four terminase subunits after Focus-refinement, colored by resolution. For each terminase subunit, the gold-standard FSC of 0.143 resolution graph and the viewing direction distribution plot are shown at middle and right, respectively.

**Fig. S4.**
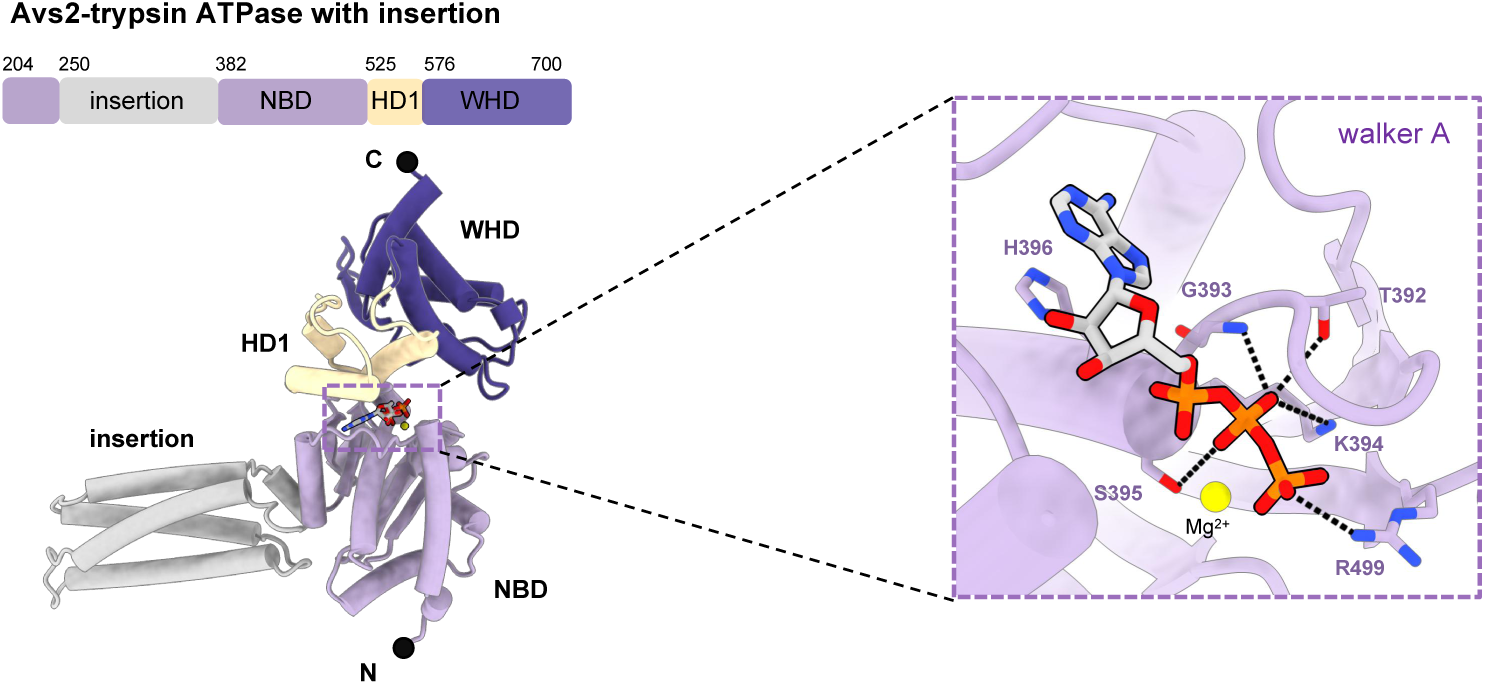
The recognition of ATP by the STAND ATPase domain of Avs2-trypsin. The cartoon representation of the STAND ATPase domain of Avs2-trypsin with the subdomains colored corresponding to the schematic diagram (top).The inset highlights the recognition of ATP by the walker A motif in the nucleotide-binding (NBD) subdomain.

**Fig. S5.**
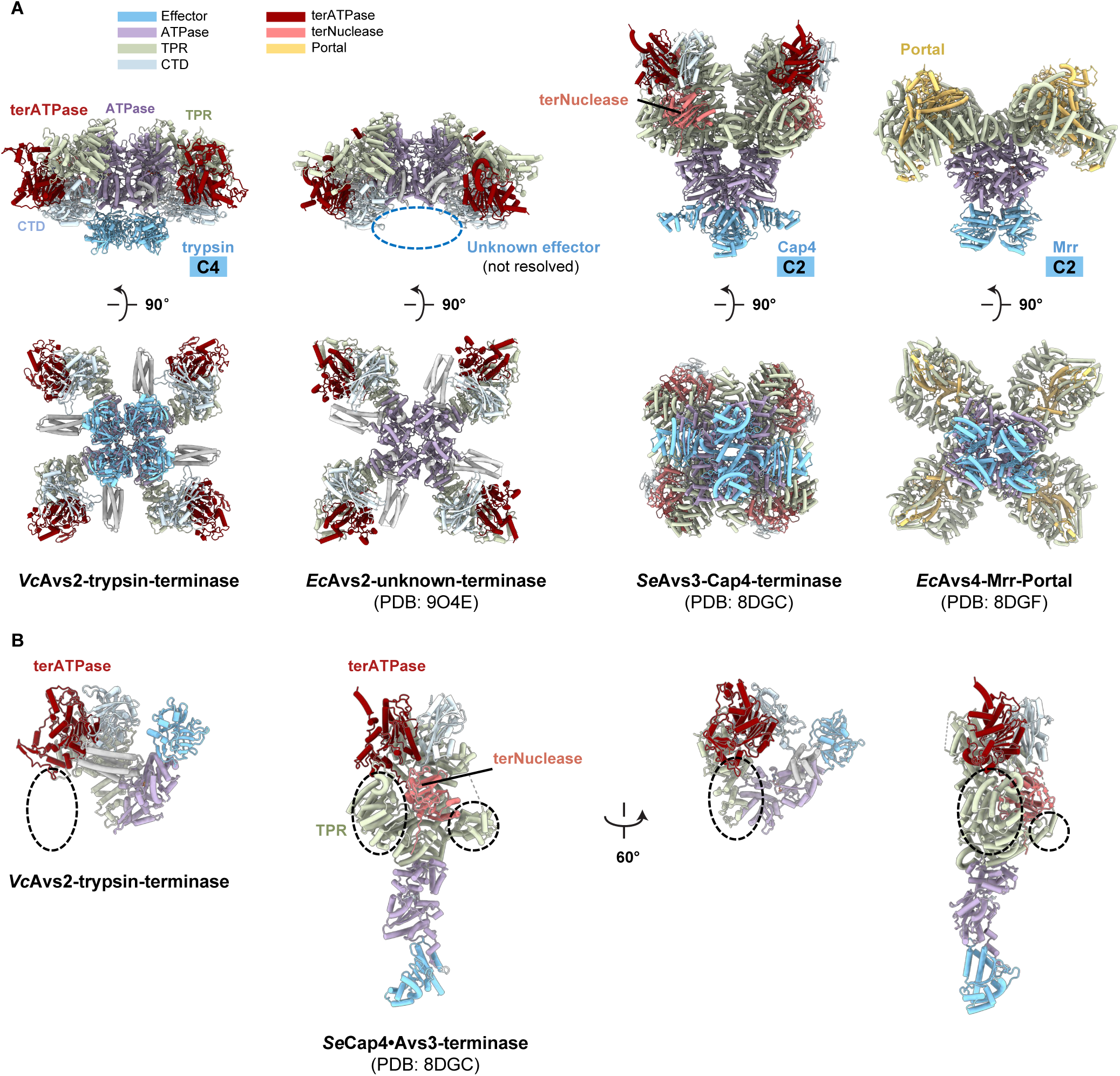
Structure comparison of VcAvs2-trypsin with its homolog and other types of Avs family. (A) Structure comparison of *Vc*Avs2-trypsin-termianse with *Ec*Avs2-unknown-terminase (PDB: 9O4E), *Se*Avs3-Cap4-terminase (PDB: 8DGC), and *Ec*Avs4-Mrr-portal (PDB: 8DGF) shows the distinctions in the overall conformation and the assembly of the effector domains. (B) Structure comparison of *Vc*Avs2-trypsin-terminase and *Se*Avs3-Cap4-terminase (PDB: 8DGC) indicates difference in TPR.

**Fig. S6.**
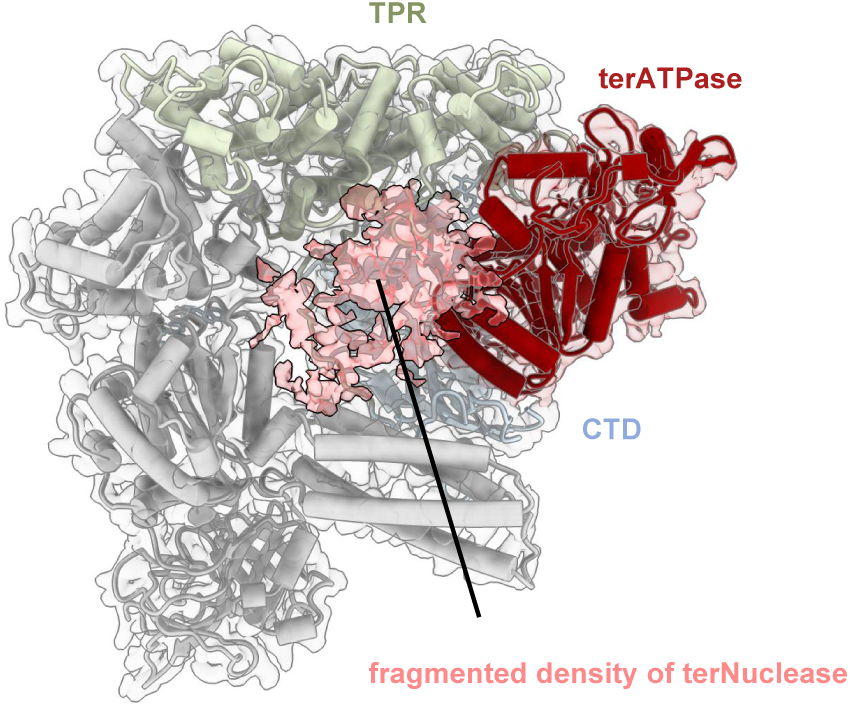
Flexibility of terminase nuclease domain. The fragmented density of terNuclease reveals its extremely flexibility due to the weak binding and recognition by Avs2.

**Fig. S7.**
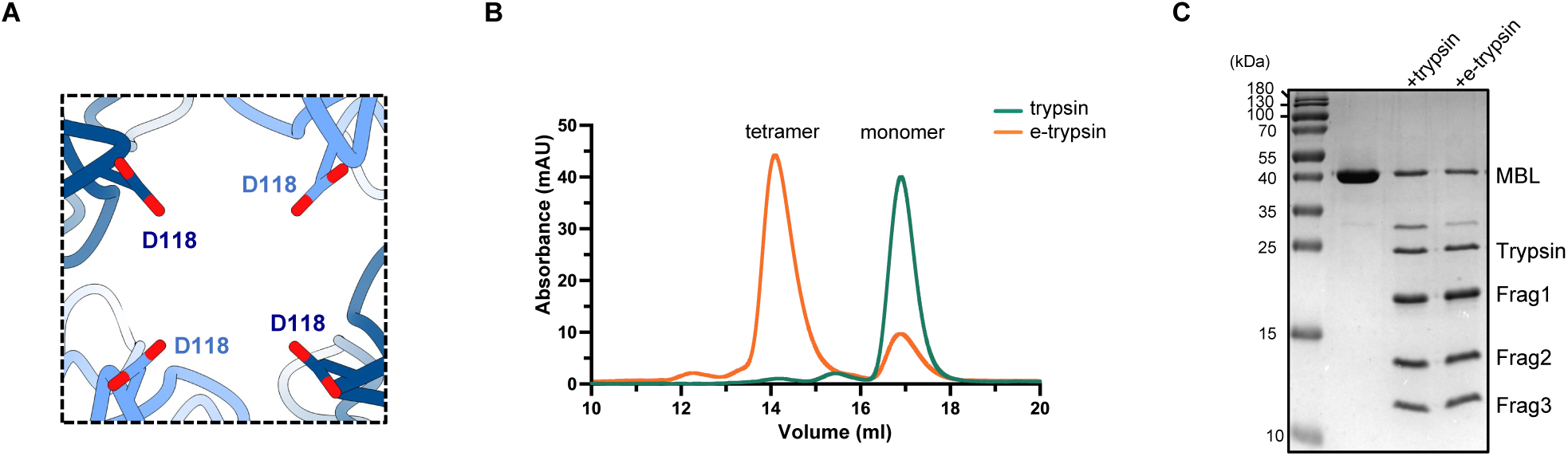
The e-trypsin domain displays enhanced ability in tetramer assembling. (A) The residue D118 at the central of trypsin tetramer interface mediates electrostatic repulsion. (B) Size-exclusion chromatography profile of trypsin and e-trypsin. (C) SDS-PAGE analysis of MBL cleavage by trypsin domain and e-trypsin. e-trypsin is a trypsin variant designed for enhanced tetramerization capacity.

**Fig. S8.**
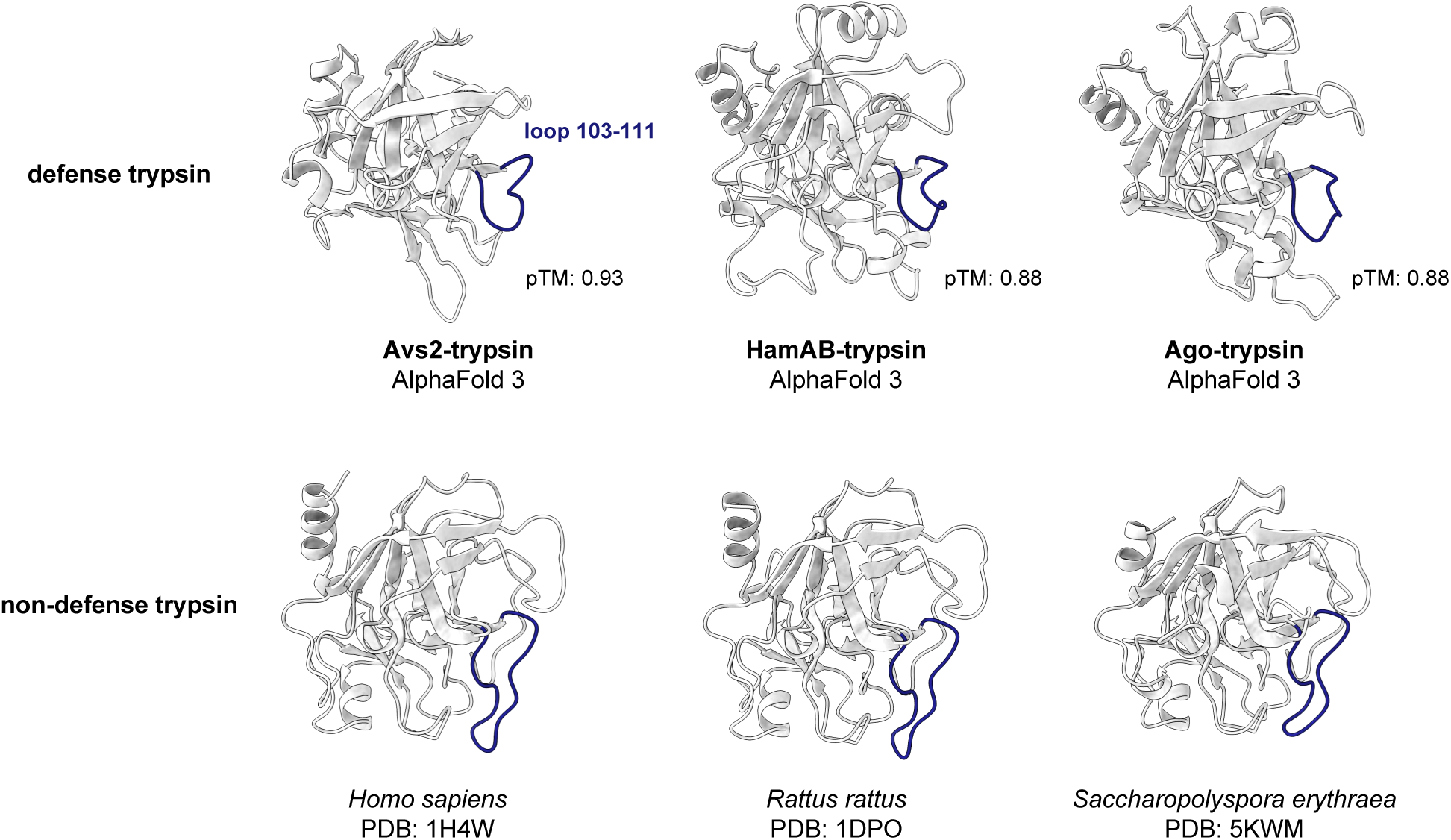
Conservation and structural divergence of the loop 103–111 in trypsin proteins. Structural comparison of trypsin domains across trypsin-MBL systems, associated with distinct sensory proteins (Avs2, HamAB, and Ago), together with representative conventional trypsin structures (PDB: 1H4W, 1DPO, and 5KWM). Loop 103–111 and its structurally equivalent regions (colored in dark blue) exhibit both short and long types, distinguishing defense from non-defense trypsin proteins.

**Fig. S9.**
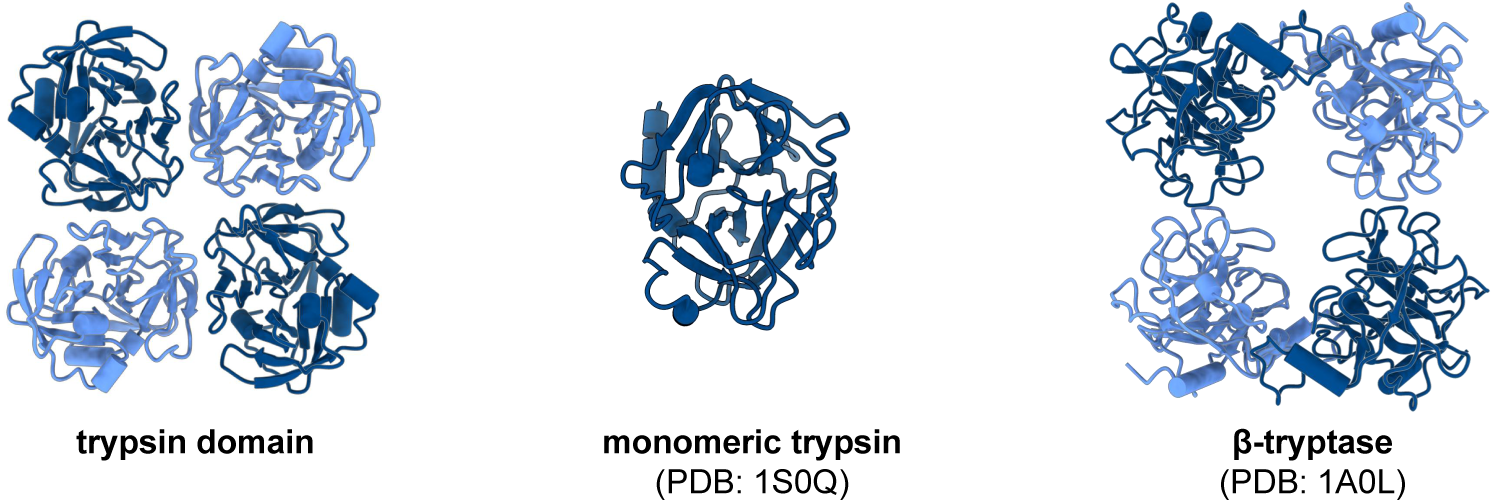
Different assemblies of trypsin proteins. Structural comparison of the active-state assemblies of the trypsin domain (left) with non-defense trypsin protein (middle, PDB: 1S0Q), and β-tryptase (right, PDB: 1A0L).

