## Supplementary Table for "Avs2 drives non-canonical tetrameric assembly of trypsin-like domain for anti-phage defense"

**Table S1 Crystallographic data collection and refinement statistics.**

| PDB ID | Cleaved-loop MBL  44AF |
| --- | --- |
| **Data collection** |  |
| Beam Line | BL18U1,SSRF |
| Space group | P2_1_ 2_1_2_1_ |
| Cell dimensions |  |
| *a, b, c* (Å) | 49.327 67.799 94.087 |
| α, β, γ (°) | 90 90 90 |
| wavelengh | 0.978610 |
| Resolution Limits (Å) | 55.01- 2.7 (2.78-2.7) |
| No.Unique reflections | 9054 (743) |
| Completeness (%) | 99.56 (99.64) |
| CC1/2 | 0.987 (0.875) |
| R-merge | 0.08376 (0.2789) |
| R-meas | 0.1185 (0.3944) |
| R-pim | 0.08376 (0.2789) |
| Mean I/sigma (I) | 6.24 (2.13) |
| **Refinement** |  |
| No.reflections | 9095 (741) |
| R-work | 0.2435 (0.3398) |
| R-free | 0.2866 (0.3823) |
| RMS (bonds) | 0.002 |
| RMS (angles) | 0.45 |
| Number of non-hydrogen atoms | 2649 |
| Macromolecules | 2638 |
| Ligands | 1 |
| Solvent | 10 |
| B-factor |  |
| Average | 26.7 |
| Ligands | 64.05 |
| **Ramachandran** |  |
| Favored (%) | 98.76 |
| Allowed (%) | 1.24 |
| Outliers (%) | 0.00 |

**Table S2 Cryo-EM data collection, refinement and validation statistics.**

| PDB ID  EMDB ID | Avs2-trypsin-terminase  27UW  EMD-81457 |
| --- | --- |
| **Data collection and processing** |  |
| Microscope | Krios G3i |
| Voltage (kV) | 300 |
| Camera | Falcon 4 |
| Magnification | 130 k |
| Pixel size (Å) | 0.932 |
| Electron exposure (e–/Å^2^) | 40 |
| Defocus range (μm) | -0.8 to -1.8 |
| Automation software | EPU |
| Symmetry imposed | C4 |
| Initial particle (no.) | 1,846,248 |
| Final particle (no.) | 178,923 |
| Map resolution (Å)  FSC threshold | 2.5  0.143 |
| **Refinement** |  |
| Atomic modeling refinement package | Phenix |
| Initial model used (PDB code) | - |
| Model resolution (Å)  FSC threshold | 2.5 |
|  | 0.5 |
| Model composition  Non-hydrogen atoms  Protein residues  Nucleotide residues  Ligands |  |
|  | 58496 |
|  | 7112 |
|  | 0 |
|  | 20 |
| *B* factors (mean, Å^2^)  Protein  Nucleotide  Ligand |  |
|  | 96.21 |
|  | - |
|  | 84.21 |
| R.m.s. deviations  Bond lengths (Å)  Bond angles (°) |  |
|  | 0.005 |
|  | 0.579 |
| Validation  MolProbity score  Clashscore  Poor rotamers (%) |  |
|  | 1.65 |
|  | 5.42 |
|  | 0.02 |
| Ramachandran plot  Favored (%)  Allowed (%)  Disallowed (%) |  |
|  | 94.88 |
|  | 5.00 |
|  | 0.11 |
| EMDB accession code | EMD-81210/EMD-81216/EMD-81451/EMD-81453/EMD-81456 |

EMDB codes are shown: the composite map, the consensus map, focused maps. Map resolution is reported for the consensus map. The model validation is reported for the composite map.
